# Somatic mutation as an explanation for epigenetic aging

**DOI:** 10.1101/2023.12.08.569638

**Authors:** Zane Koch, Adam Li, Daniel S. Evans, Steven Cummings, Trey Ideker

**Affiliations:** Program in Bioinformatics and Systems Biology, University of California San Diego, La Jolla CA, 92093, USA; California Pacific Medical Center Research Institute, San Francisco CA 94158, USA; Department of Epidemiology and Biostatistics, University of California, San Francisco, CA, 94158; Department of Medicine, University of California San Diego, La Jolla California, 92093, USA

## Abstract

DNA methylation marks have recently been used to build models known as “epigenetic clocks” which predict calendar age. As methylation of cytosine promotes C-to-T mutations, we hypothesized that the methylation changes observed with age should reflect the accrual of somatic mutations, and the two should yield analogous aging estimates. In analysis of multimodal data from 9,331 human individuals, we find that CpG mutations indeed coincide with changes in methylation, not only at the mutated site but also with pervasive remodeling of the methylome out to ±10 kilobases. This one-to-many mapping enables mutation-based predictions of age that agree with epigenetic clocks, including which individuals are aging faster or slower than expected. Moreover, genomic loci where mutations accumulate with age also tend to have methylation patterns that are especially predictive of age. These results suggest a close coupling between the accumulation of sporadic somatic mutations and the widespread changes in methylation observed over the course of life.

## Introduction

Practically since the elucidation of the DNA double helix, it has been postulated that progressive damage to this fundamental structure is the cause of aging^1–5^. The primary support for this theory relates to somatic mutations, which accumulate in the genomes of most tissues and species throughout life^4,6–9^. Such accumulation has been associated with multiple characteristics of old age, including immune dysfunction^10–12^, neurodegeneration^13–15^, and cancer^16–19^.

Aging has also been associated with other major types of molecular changes beyond DNA mutations^20,21^, leading to debate as to which of these aging “hallmarks” are fundamental causes^22–25^. In particular, much recent attention has been given to associations of age with DNA methylation, a dynamic epigenetic mark found primarily at CG dinucleotides (CpG sites) throughout the genome^26^. CpG methylation has diverse functional consequences including X chromosome inactivation^27,28^, chromatin and transcriptional regulation^26,29^, cell-type specification, and maintenance of pluripotency^30–32^. DNA methylation patterns have been found to change very regularly over the course of life, prompting the creation of statistical models, termed ‘epigenetic clocks’, which attempt to predict an individual’s age using their DNA methylation profile^33–35^. Subsequent research has shown that epigenetic clock predictions correlate with a host of age-related biological attributes, including frailty, Alzheimer’s disease, all-cause mortality, life-extending intervention^36^, and time-to-death^37–40^. Such observations have bolstered epigenetic theories of aging, which propose that progressive remodeling of the epigenome leads to aging phenotypes via the dysregulation of gene expression, cellular function, and senescence^24,41–43^. The degree to which epigenetic changes are direct causes of aging, however, remains unclear.

Despite the separate interest in DNA mutations and DNA methylation as theories of aging, the relationship between the two processes is not well understood. One recent study reported that somatic mutations in DNA-binding sites for Tet Methylcytosine Dioxygenase 1 (TET1), the primary enzyme involved in the removal of methylation marks^44^, are associated with local hypermethylation^45^. Another study demonstrated an association between somatic mutations, subsequent clonal expansion of blood cells, and accelerated epigenetic aging^46^. Most other research linking DNA sequence and methylation has focused on inherited germline variants rather than acquired somatic mutations, such as efforts to identify methyl quantitative trait loci (me-QTLs) linking common polymorphisms to methylation levels of specific CpG sites^47–49^.

Nonetheless, an intrinsic biochemical connection between DNA mutation and methylation occurs at 5-methyl cytosine residues^6,50,51^, which spontaneously deaminate over time to yield thymine^52^. A prerequisite for this mutational event is cytosine methylation, relating somatic mutation to prior epigenetic modification of DNA. Conversely, a prerequisite for DNA methylation is the presence of a cytosine, which may be eliminated by prior somatic mutation. Given this interdependence, we considered that the separate links that have been established between DNA mutation and aging, and DNA methylation and aging, might each reflect a common underlying process whereby methylation potentiates mutation and/or mutation potentiates changes in methylation.

To explore this hypothesis, we set out to comprehensively examine the relationship between somatic DNA mutations and DNA methylation in large collections of human tissue samples characterized for both layers of molecular information. In what follows, we identify several types of interaction between somatic mutation and DNA methylation, both one-to-one and one-to-many (**Fig. 1a**). Based on these findings, we use somatic mutations as a surrogate for epigenetic marks in measures of aging, indicating the degree to which epigenetic aging is explained by somatic mutations (**Fig. 1b**).

**Figure 1:**
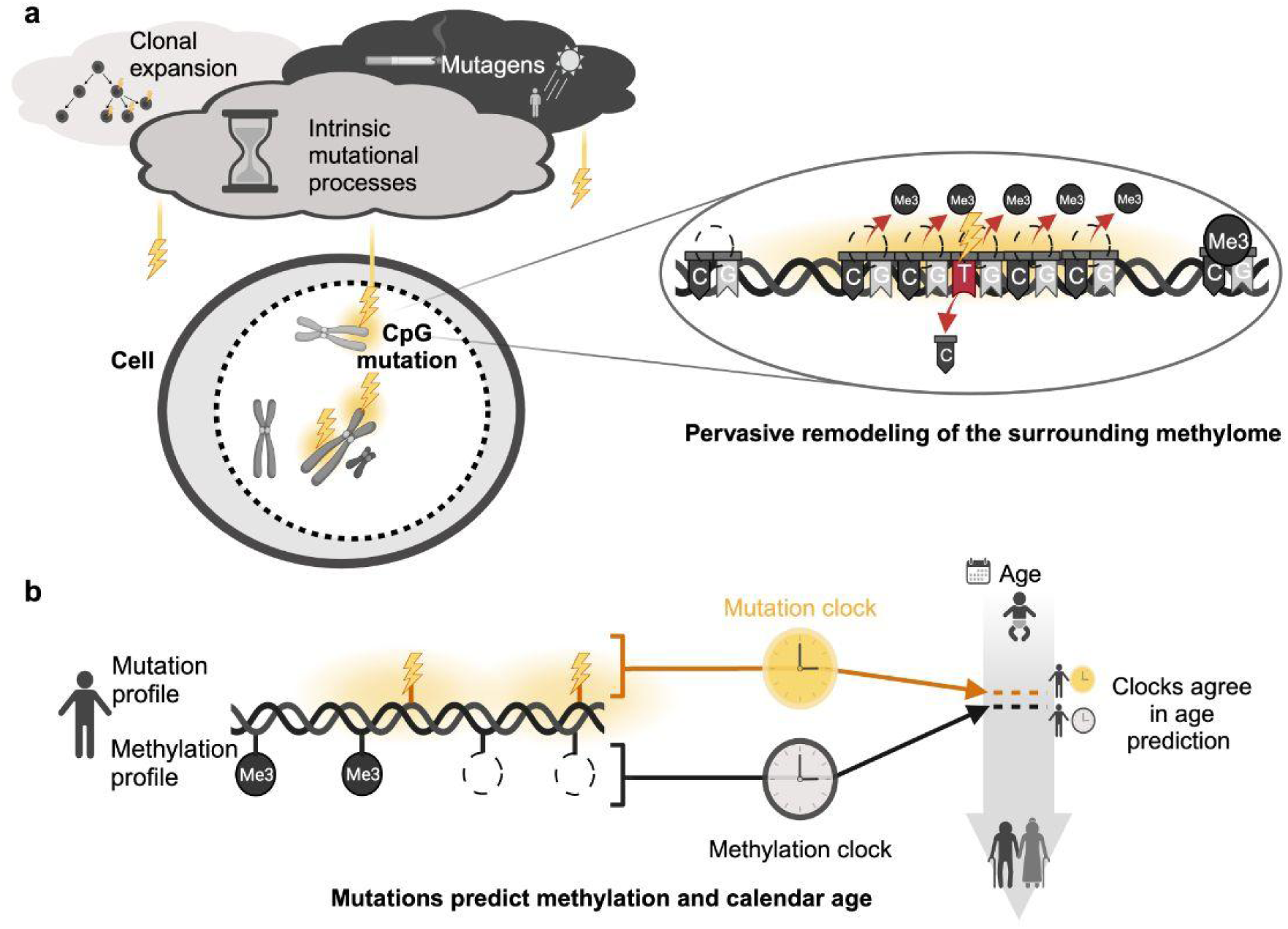
Links among CpG mutations, methylome remodeling, and aging. **a)** Various mutational processes affect the genome. Here, we show that some of these mutations associate with an aberrant DNA methylation pattern at both the mutated site and at numerous neighboring CpGs. **b)** An individual’s DNA mutation profile and DNA methylation profile make similar predictions of both their calendar age and rate of aging.

## Results

### Genome-wide hypomethylation of mutated CpG sites

To study the connections between somatic mutations and DNA methylation marks, we analyzed multi-omics data from human patients cataloged in The Cancer Genome Atlas (TCGA)^53–55^ and the Pan-Cancer Analysis of Whole Genomes (PCAWG)^56^. Tumor biopsies had been drawn from a diversity of tissue types and characterized by whole-exome sequencing (TCGA, 8,680 exomes across 33 tissues) or whole-genome sequencing (PCAWG, 651 genomes across 7 tissues). In each case, DNA from the tumor sample was compared to a second DNA sample drawn from the same individual, with differences used to define somatic mutations (typically comparing the tumor DNA sequence to whole blood, **Methods**). These data had been complemented by methylation profiling of the same tissues via the Illumina Infinium HumanMethylation450 BeadChip, which provides methylation fraction readouts (the fraction of DNA reads that are methylated) for approximately 450,000 CpG sites genome-wide^57^.

From these data, we considered all single base-pair substitution mutations (n = 3,457,875 mutation events) and CpG sites for which all individuals had a reliably measured methylation value (n = 326,751 CpG sites, **Methods**). Consistent with previous reports^6,50^, CpG sites were the most frequently mutated dinucleotide accounting for 13.5% of all somatic mutations genome-wide (**Fig. 2a**). The vast majority of these were C>T transitions (82.3%, **Supplementary Fig. 1a**) occurring at sites that tended to be heavily methylated (**Supplementary Fig. 1b**).

**Figure 2:**
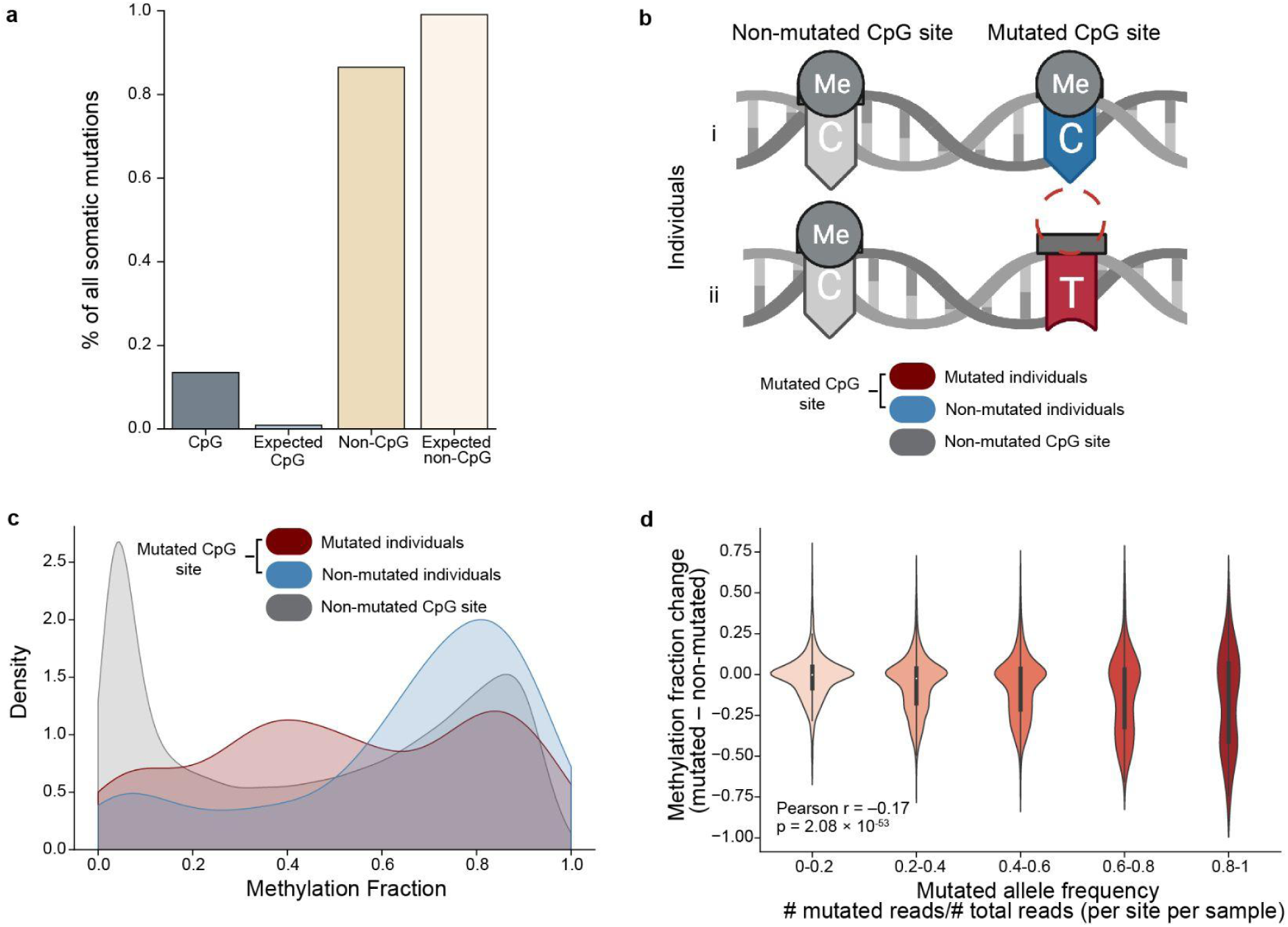
Frequency and methylation status of CpG mutation events. **a)** Percent of genome-wide somatic mutations that are classified as CpG (n = 467,079 mutations) or non-CpG mutations (n = 2,990,796 mutations). Expected percentages are calculated supposing mutation probability to be uniform across the genome (**Methods**). **b)** Diagram showing two categories of CpG sites: those where no individual is mutated (non-mutated CpG site, gray) and those where a mutation has occurred in at least one individual (mutated CpG site, red) and remaining individuals are non-mutated (blue). **c)** Distribution of CpG methylation values for the categories of CpG sites from (b). The methylation fractions of mutated individuals (red) and non-mutated individuals (blue) are shown for the 1,000 CpG sites with the highest mutated allele frequency (corresponding to MAF > 0.53, **Methods**). **d)** Methylation change between mutated and non-mutated individuals at (n = 8,037) mutated CpG sites. Methylation change is the difference between the median methylation fraction in mutated individuals and the median methylation fraction in non-mutated individuals of matched age and tissue. CpG sites are binned into five groups based on MAF, with violin plots summarizing the distribution of methylation changes within each group. Vertical bars inside each violin represent the interquartile range.

We next asked whether individuals harboring a mutated CpG site exhibit lower levels of methylation at that site compared to non-mutated individuals (**Fig. 2b**). We reasoned that once mutated, the site would no longer constitute a CG dinucleotide, reducing its likelihood of methylation. Indeed, we found a significant decrease in methylation in individuals with a mutation at a CpG site compared to non-mutated individuals at the same site (Mann-Whitney p = 3.90⨉10^−9^, **Fig. 2c, Methods**), with loss of methylation proportional to the frequency of reads with the mutant allele (Pearson r = –0.17, p = 2.08⨉10^−53^, **Fig. 2d**). These results supported a model in which CpG mutations occur primarily at hypermethylated sites due to the spontaneous deamination of methylcytosine and can become fixed in the genome of daughter cells causing a decrease in methylation proportional to the mutated clonal population.

### Mutated sites show extensive remodeling of the surrounding methylome

During this exploration, we noted numerous cases in which somatic mutations coincided not only with hypomethylation at the mutated CpG site but also with atypical methylation of numerous CpGs in the surrounding genome. An illustrative example was the C>T mutation at base pair 56,642,556 of chromosome 16 in the individual TCGA-GV-A3QI (**Fig. 3a**). CpG sites adjacent to this somatic mutation were strikingly hypermethylated in this individual, with such hypermethylation extending over a contiguous region more than 30 kb downstream. This effect encompassed the metallothionein 2A gene as well as additional metallothionein family members MT1E and MT1M, for which methylation-linked repression has been associated with metastasis in multiple cancer types^58–61^.

**Figure 3:**
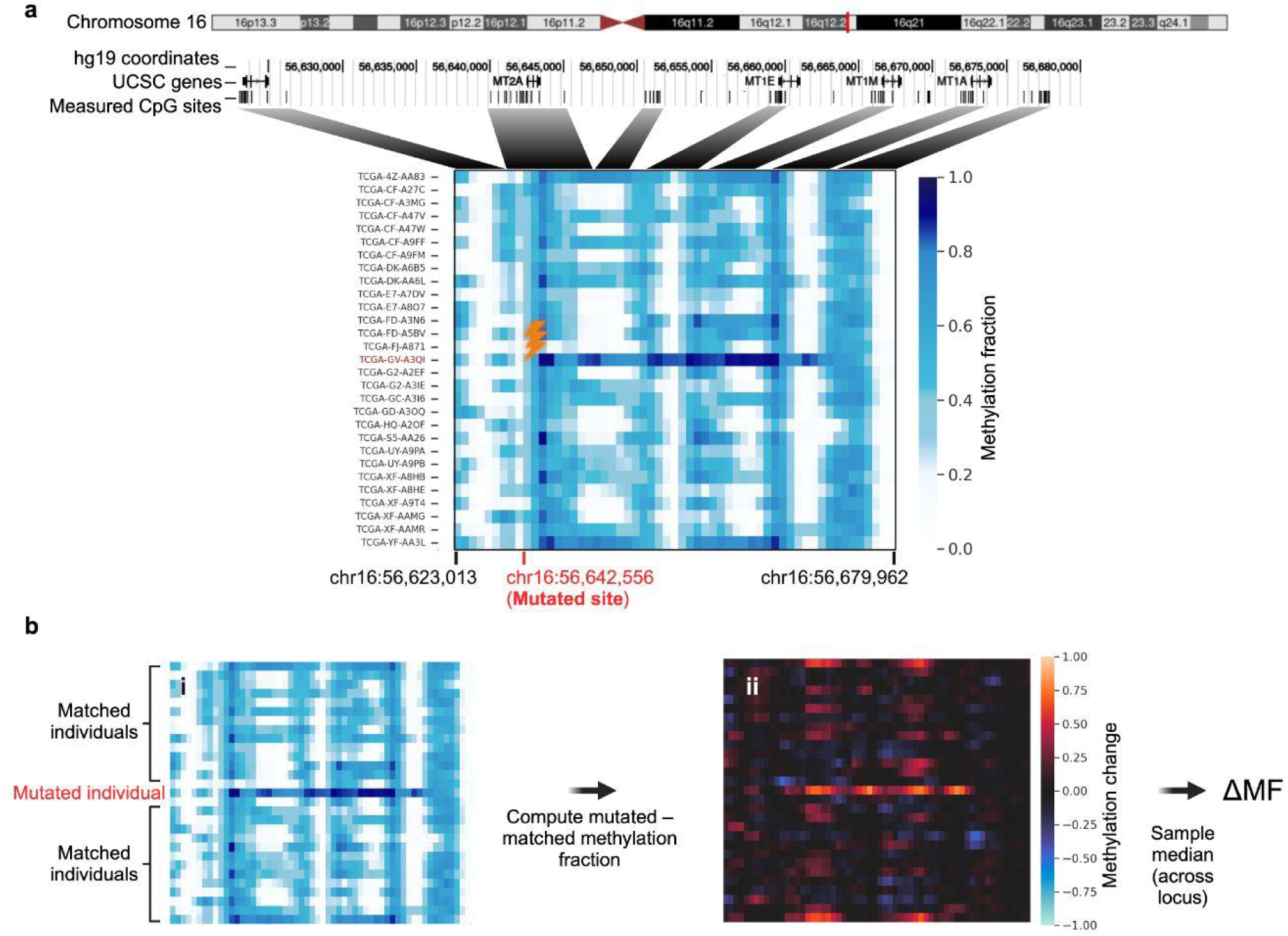
Association of mutations with regional methylation patterns. **a)** Example mutated site where the individual TCGA-GV-A3QI has a C>T mutation at chr16:56,642,556 of the hg19 human genome. **Upper**: Ideogram of chromosome 16, with a red bar indicating the location of the mutated site. The first underlying track shows hg19 base pair coordinates, the second the documented genes in the region, encoding five Metallothionein (MT) factors, and the third the locations of CpG sites measured on the Illumina 450k methylation array (vertical bars). **Lower**: Heatmap of CpG methylation fractions. Rows are samples (1 mutated, 28 background), and columns are the measured CpGs within a ±50 kb window proximal to the mutation (n = 62 CpG sites). The color corresponds to the methylation fraction of each CpG. The mutated sample row and mutated site column are labeled in red, with the mutation event indicated by a lightning bolt. **b)** Calculation of change in methylation fraction, or ΔMF, with reference to a specific mutated site. **i)** Heatmap of methylation fractions of the mutated site and CpGs in the surrounding window, replicated from panel (a). **ii)** Heatmap of corresponding differences in methylation between each sample (row) and all other samples in the matrix (median of other rows), computed separately for each site in the window (columns). The final ΔMF value was calculated as the overall methylation change of the mutated sample, taking the median across all sites in the window (**Methods**).

To move beyond anecdotal observations, we devised a general test for whether somatic mutations are associated with remodeled methylation at surrounding CpGs. In a window centered on each mutated site, we computed a quantity we called ΔMF: the change in methylation fraction observed for CpGs in the window, comparing the mutated individual to matched non-mutated individuals (**Fig. 3b, Methods**). We observed that ΔMF tended toward substantially more extreme values than expected at random (n = 2,600 mutated sites with sufficient nearby CpGs, p < 10^−124^, **Fig. 4a**), with mutated loci more than four times as likely to have an extreme decrease in nearby methylation (ΔMF < –0.3, **Fig. 4b**). Mutated loci were also enriched for nearby methylation increases, albeit more weakly (**Fig. 4b**). Examination of different window sizes showed that the methylation increases/decreases were localized to ±10 kb from the site of mutation, with CpGs close to the mutated site having the most extreme methylation changes (**Fig. 4c**). Deeper explorations revealed that the aberrant methylation patterns: [1] are specific to genomic context, occurring exclusively at mutations to CpG sites (**Fig. 4d**); [2] have a direction of change that depends on local CpG density (i.e., whether they are inside CpG islands^29^, **Fig. 4d**); and [3] increase proportionally with the fraction of DNA in the sample harboring the mutation (**Fig. 4e**). These results indicated that our earlier observations were not anecdotal but that CpG mutations are generally associated with an atypical pattern of methylation in the surrounding DNA.

**Figure 4:**
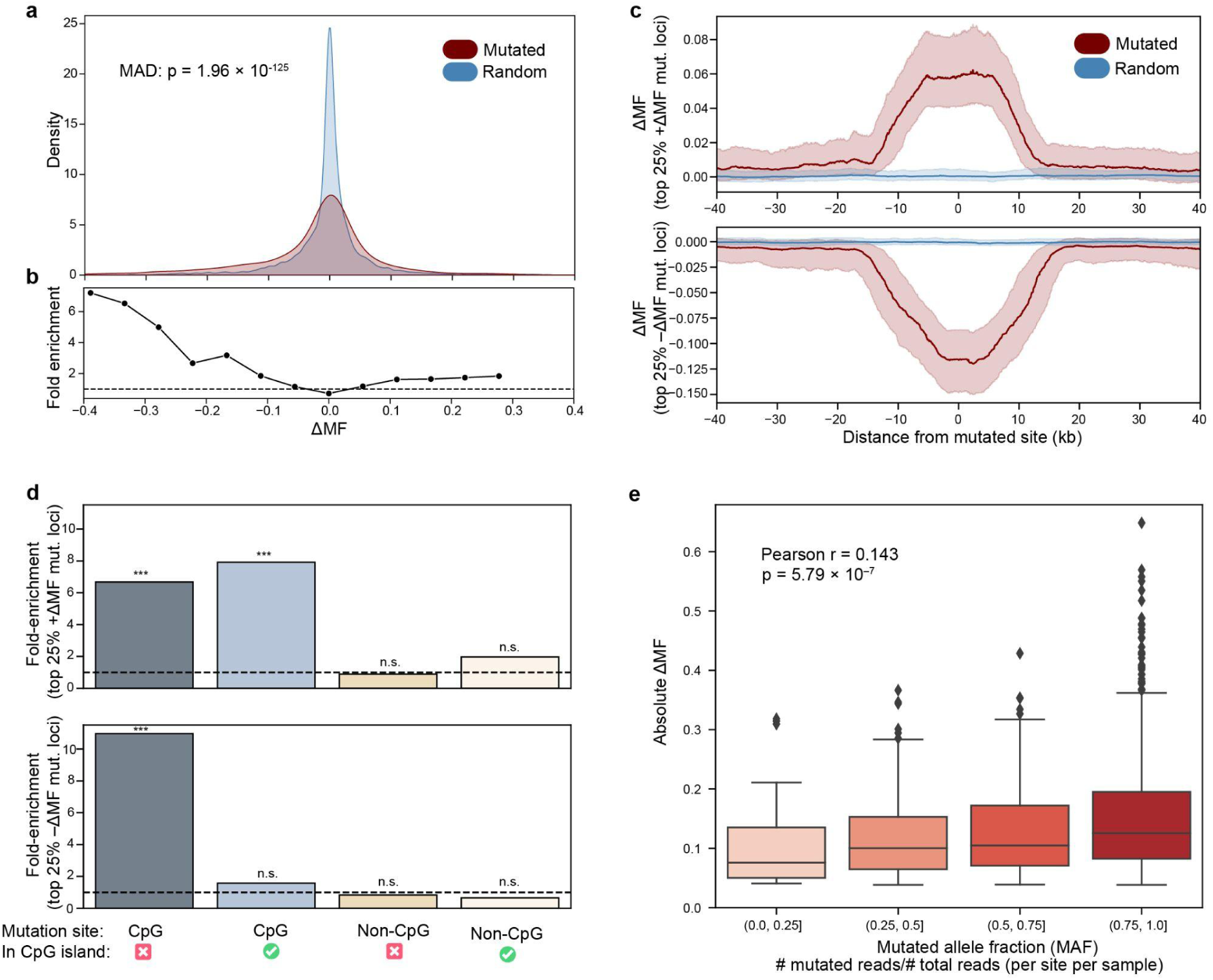
Magnitude and extent of methylation changes near somatic mutations. **a)** Probability distribution of ΔMF values calculated in a ±10 kb window surrounding mutated (red) versus random control (blue) sites. Mutated sites include n = 2,600 mutated sites with MAF ≥0.8, ≥15 matched individuals (individuals of same tissue type within ±5 years of age), and ≥1 measured CpG within the window. Random control sites include n = 260,000 non-mutated sites (**Methods**). P value shown for a two-sided Mann-Whitney test for a difference in median absolute deviation (MAD) of ΔMF between the mutated and non-mutated random control loci. **b)** Line plot depicting the fold enrichment for mutated over non-mutated sites as a function of ΔMF. Fold enrichment is the ratio of the probability of observing a given ΔMF for mutated sites versus the probability of that ΔMF for non-mutated control sites. ΔMF divided into equally spaced bins from –0.4 to 0.4. **c)** Line plot depicting ΔMF as a function of genomic distance from the site of mutation. For the 25% of mutated sites with the most positive (**top**, n = 650) or negative (**bottom**, n = 650) ΔMF values from (a), the ΔMF value in overlapping 2 kb windows at each distance from the mutation is plotted for mutated sites (**red**) versus random control sites (**blue**). The shaded region indicates the 40th-60th percentiles at that same distance. **d)** Enrichment of extreme ΔMF values at CpG sites and CpG islands. Top versus bottom barcharts show the 25% of mutations with the most positive versus most negative ΔMF values in panel (a) (n = 650 mutations each). The enrichment of these mutations (bars, y axis) is considered for different types of sites, depending on whether the site is a CpG and/or falls within a CpG island (x-axis categories). Enrichment is compared to the genomic baseline (**Methods**), with significance determined by a one-sided binomial test. Significant enrichment (p ≤ 0.001) is marked with (***), and non-significant (p > 0.01) is marked with (n.s.). CpG Islands are defined as genomic regions ≥ 200 bp, ≥ 50% GC content, and a high CpG occurrence. **e)** Boxplot of the absolute ΔMF value as a function of the mutated allele fraction (MAF). Includes all mutated sites with ≥15 matched samples (samples of the same tissue type within ± 5 years of age) and ≥1 measured CpG within ±10 kb (n = 3,880 mutated loci). Two-sided p value calculated based on the exact distribution of Pearson’s r modeled as a beta function.

### Somatic mutations mirror epigenetic predictions of age

While mutation of any particular CpG site is exceedingly rare in the human population and thus a poor predictor of age, its corresponding CpG methylation fraction varies regularly in a manner that is often age-associated^33^. We considered, however, that the one-to-many relationship revealed by our previous analysis (**Fig. 4**), by which a single CpG mutation maps to a broad profile of methylation changes in the surrounding DNA, might bridge this apparent gap between sporadic mutation accumulation and consistent methylation change. Accordingly, we compared two procedures for the prediction of human chronological age: the first using an individual’s profile of CpG methylation values, as in previous epigenetic aging models (methylation clock), and the second using their profile of somatic mutations, including the counts of somatic mutations within 10 kb of each of these same CpGs (mutation clock, **Methods, Fig. 5a**). Evaluating these models using a nested cross-validation procedure (**Methods**), we found that the methylation clock predicted age with an accuracy of r = 0.77 (Pearson correlation), while the mutation clock had an accuracy of r = 0.70 (**Fig. 5b-c**). Examining predictions within each individual tissue, both models were most accurate at predicting age in brain samples and least accurate in thymus samples (**Supplementary Fig. 2**).

**Figure 5:**
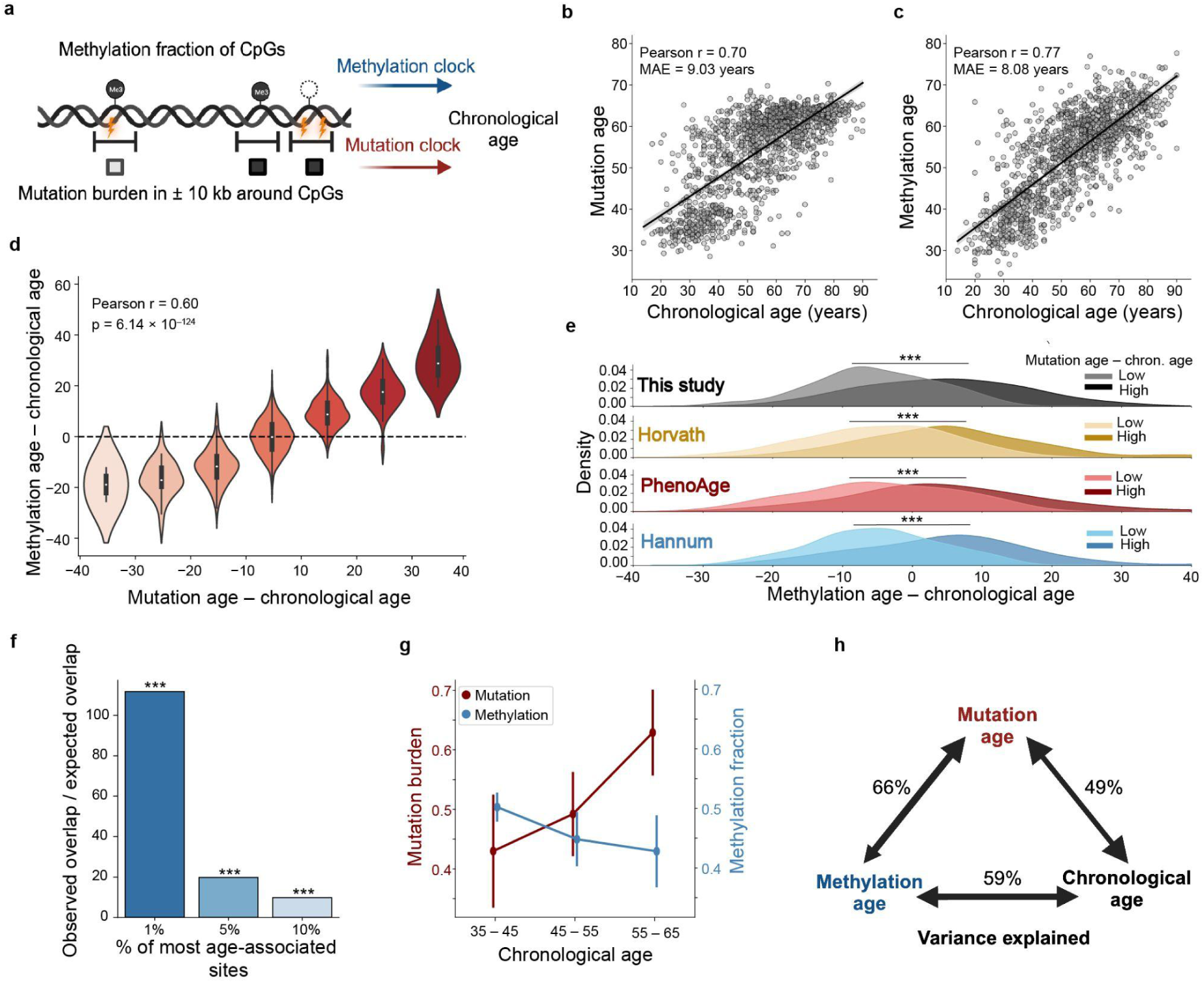
Association between mutation age, methylation age, and chronological age. **a)** Methylation clock: the methylation fractions of CpGs are used in a gradient boosted tree model to predict chronological age. Mutation clock: the count of mutations around the same CpGs is used in an identical model to predict chronological age. Both models incorporate similar covariates (**Methods**). **b)** Scatter plot of human individuals, showing age predictions from the mutation model versus their chronological age. Includes 1,250 individuals from five tissues (**Methods**). **c)** Similar to panel (b) but showing age predictions from the methylation rather than mutation model. **d)** Violin plots of the methylation age residual versus mutation age residual. The residual in each case is the predicted age minus chronological age. Plot includes the same individuals as in panels (b,c). Pearson r refers to the correlation between methylation age and mutation age, controlling for chronological age (i.e., partial correlation, p = 6.14 10^−124^). **e)** Distribution of methylation age residuals for the same individuals as in panels (b,c), computed according to each of four previous methylation clocks. “This study” refers to the methylation clock shown in panel (c) (**Methods**). For each clock, the 20% (n = 250) of individuals with the youngest mutation age for their chronological age are shown in lighter color (low mutation – chronological age), and the 20% (n = 250) of individuals with the oldest mutation age for their chronological age are shown in darker color (high mutation – chronological age). (***) indicates a significant (p ≤ 10^−10^) difference in distribution between the low and high mutation residual age groups, based on a two-sided Mann–Whitney U test. **f)** Barplot depicting the ratio of observed to expected overlap between sets of age-associated CpG sites. The CpGs with maximal (top 1%, 5%, and 10%) mutual information between local mutation burden (±10 kb) and age or between methylation fraction and age were chosen. The intersection (overlap) between age-associated mutation burden and age-associated methylation sets was compared to the expected intersection assuming random selection (**Methods**). Significant enrichment (p ≤ 10^−10^) is marked with (***). **g)** Mutation burden (y-axis left) or methylation fraction (y-axis right) is plotted versus chronological age (x-axis) for CpG site cg19236454. Data from brain (LGG) samples, considering individuals with a nonzero mutation burden (±10 kb) at this site (n = 67). Pearson correlation with chronological age: mutation burden = 0.18, methylation = –0.18. Error bars denote standard error. **h)** Diagram summarizing the relationships between three measures of age: mutation, methylation, and chronological time. Variance explained is calculated as the squared Pearson correlation between each pair of measures for the same individuals as in panels (b,c).

Beyond their similar accuracies of age prediction, we found that the two clocks agreed significantly in several other key aspects. First, the predictions from both models were highly correlated across individuals (r = 0.81), and this relationship persisted even after controlling for calendar age (partial correlation = 0.60, p = 6.14⨉10^−124^, **Fig. 5d**, **Methods**). For example, for individuals predicted by mutations to be one year older than their calendar age, the methylation clock yielded a corresponding overprediction of 0.75 ± 0.53 years (mean ± stdev). This same agreement in over/underprediction (similarity in model residuals) was observed when comparing the mutation clock to previously published methylation clocks (**Fig. 5e**)^33,34,39^. Second, CpG sites for which the surrounding mutation burden was most associated with age also tended to have the most age-associated methylation values (**Fig. 5f**, **Methods**). One example was the CpG site cg19236454 (chr19:42,799,926), for which the local mutation burden progressively increased with age (±10 kb, r = 0.18), while the methylation of the site was progressively lost (r = –0.18, **Fig. 5g**). Thus, mutation and methylation profiles were synchronized with respect to predictions of age (**Fig. 5h**), both globally (genome-wide) and locally surrounding individual CpG sites.

## Discussion

In this study, we have observed notable associations between CpG mutation and methylation at multiple scales. At the scale of single nucleotides, CpG sites altered by somatic mutation, the most frequent mutation type genome-wide (**Fig. 2a**, **Supplementary Fig. 1a-b**), exhibit loss of methylation at that site (**Fig. 2c-d**). At a larger scale, such mutated sites coincide with sweeping methylation changes across numerous CpGs within the surrounding genomic region (**Fig. 3**, **4a-c**). Plausibly as a result of this larger-scale relationship, individuals whose mutations indicate increased genomic age also tend to have older methylomes (**Fig. 5d, h**).

A fundamental tension addressed in this study is that two individuals very rarely share a somatic mutation at the same CpG site; thus, mutations would initially seem too sparse to explain the numerous CpG sites at which methylation reliably changes with age. However, our findings show that single mutations can correspond to appreciable shifts in the methylome, with a graded relationship that depends on the frequency of the mutated allele (i.e., clonality of the mutant cell population, **Fig. 4e**). Consistent with these findings, we see that within individuals of the same calendar age, mutation and methylation clocks agree on which individuals are aging faster or slower (**Fig. 5d-e**) and that somatic mutations explain almost 50% of the variation in methylation age across individuals (**Fig. 5b, h**).

The mechanisms by which a CpG mutation affects its methylation state or, conversely, CpG methylation potentiates its own mutation, have already been established. The prior methylation of a CpG makes a subsequent somatic mutation more likely due to methylcytosine deamination^52^. In turn, when either nucleotide of the CpG site is mutated, the site is no longer a CpG, substantially decreasing the likelihood of future methylation by a DNA methyltransferase^62^.

For mutations exhibiting larger-scale gains or losses of methylation in the surrounding kilobases, it is conceivable that either methylation or mutation, or neither, may be the primary causal agent. The observed association between somatic mutation and local hypermethylation (**Fig. 4a**) may occur if hypermethylation creates an environment prone to methylcytosine deamination events, giving rise to rare somatic mutations embedded within hypermethylated regions. This model does not, however, explain the frequently observed co-occurrences of mutations with neighboring *hypo*methylation throughout the genome (**Fig. 4a**), as hypomethylation should decrease, not increase, the local probability of mutation.

An alternative possibility is that mutations are the primary causes of subsequent changes in methylation. Mutations within the DNA binding site of a methylase or demethylase enzyme could plausibly affect enzyme activity, dysregulating the methylation state of the surrounding genome^63–65^. Such a relationship has been reported explicitly for somatic mutations in the DNA binding sites of TET1, a demethylase, leading to local gains of methylation^45^. More broadly, it is well known that germline DNA sequence variants govern the methylation patterns of many CpG sites, affecting as many as 40% of CpGs genome-wide (“meQTLs”)^66^. Somatic mutations in these sequences may yield effects on methylation analogous to those observed for inherited variants.

A third possibility is that mutation and methylation events are not causal of each other, but that both are downstream of some earlier event. One such event might be related to the repair of DNA double-strand breaks (DSBs), which have been demonstrated to result in both somatic mutations and methylation changes near the site of repair^67–72^. Here, the mutation and methylation changes would be indicative of an earlier DSB repair, an activity recently suggested to cause epigenetic aging^41^.

Regardless, understanding the causality between mutations, methylation, and aging has important implications for how we seek to prevent or reverse aging. In particular, if mutations are the fundamental driver of aging phenotypes and epigenetic changes simply track this process, then strategies aimed at epigenetic reversal^41,73–77^ may be treating a symptom rather than a cause.

As the current large human datasets measuring both somatic mutations and methylation pertain to cancer patients, the relevance of our findings to normative aging should be examined, i.e., in normal individuals and tissues. This limitation notwithstanding, we would note that mutation burdens in cancerous and normal tissues are similar^78,79^ and that the majority of mutations found in tumors are thought to represent normal mutational processes unrelated to cancer^78,80^. A second limitation is that our analysis is cross-sectional rather than longitudinal, with each individual measured at a single time point only. Here, a longitudinal study design could greatly inform the actual order of events. In addition, there exist other factors associated with epigenetic aging that do not explicitly implicate somatic mutations. Some epigenetic changes clearly reflect alterations in tissue composition with age^81,82^, and other changes are associated with the expression of developmental genes^83–89^ such as in the binding sites of the polycomb repressive complex^90,91^. Some of these factors may nonetheless relate to DNA mutations, for instance somatic mutations can drive alterations in tissue composition^92,93^.

## Online Methods

### Data access and preprocessing

We obtained paired DNA methylation (Illumina 450k array) and somatic mutation data from two public consortia: TCGA^53–55^ and PCAWG^56^. Relevant to TCGA, we used the PAN-CAN cohort (http://xena.ucsc.edu/), which includes 8,680 samples from 33 cancer types with both Illumina 450k methylation array data and somatic mutation calls. Data from the PCAWG consortium (https://dcc.icgc.org/pcawg) include 651 samples from 7 cancer types with both Illumina 450k methylation array data and whole-genome somatic mutation calls. Methylation data from both cohorts were further processed as follows. First, we removed CpG sites for which any sample had a missing value, leaving 273,202 CpG sites for TCGA and 326,749 CpG sites for PCAWG. Second, we removed samples for which the mean methylation fraction (over all remaining CpGs) was more than three standard deviations outside of its expected (mean) value over all samples. Third, each sample was quantile normalized.

### Characterizing CpG mutation frequency

Based on UCSC hg19 human genome annotations^94^, the number of nucleotides that comprise CpG residues equals 2 bp * 28,299,634 CpG sites, within a total genome length of 3,137,144,693 bp. Therefore, 1.8% of randomly distributed mutations are expected to be CpG mutations, and the remaining 98.2% of mutations are not (**Fig. 2a**). As CpG sites are palindromic, CG on one DNA strand is equivalent to GC on the complementary strand; thus, for simplicity, we refer to all CpG mutations on either strand as alterations to the C residue in the first position. This convention was used to record the frequency of each dinucleotide sequence resulting from a CpG mutation (**Supplementary Fig. 1a**). For these cumulative analyses relating to the overall frequency of CpG mutation (**Fig. 2a, Supplementary Fig. 1a-b**), the PCAWG samples were used exclusively because they had whole genome sequences, encompassing all CpG sites, rather than exome sequences only.

### Characterizing methylation at mutated CpG sites

The methylation status of two categories of CpG sites were compared: “Non-mutated sites”, where no mutation was observed in any individual, and “Mutated sites”, where at least one individual had a mutation (**Fig. 2b**). For CpG sites of the first category (265,399 non-mutated sites), the distribution of methylation fractions was plotted (**Fig. 2c**). For CpG sites of the second category (8,037 mutated sites), some individuals harbor a mutation at that particular CpG, and some do not. In this case, the distributions of CpG methylation fractions were plotted separately for the mutated versus non-mutated individuals (**Fig. 2c**). For analyses of methylation associated with mutated sites (**Fig. 2c-d**), TCGA samples were used exclusively, as there were many more occurrences of CpG mutations in this dataset due to its much larger sample size.

### Calculating mutation-associated methylation change

Somatic mutation events (*i*, *j*) were defined as (site, sample) pairs for which the nucleotide present at site *i* in sample *j* assumed a different (A,C,G,T) value than in the matched normal tissue. For each mutation event, the genomic “locus” was defined as a ±*w* kilobase window (upstream and downstream) centered on the mutated site *i*. Matched background samples *b* were defined as those without any somatic mutations in this window and that were of the same tissue type and approximate age (±5 years) as the mutated sample. The methylation fraction of CpG site *k* in sample *j* is denoted by *m_k,j_*. Using these definitions, we calculated ΔMF, the median change in methylation fraction in the window comparing sample *j* to matched background samples.

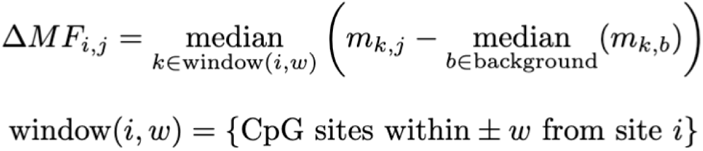

We calculated ΔMF in ±10 kb windows for each mutation event in the PCAWG data, where [1] the mutation was a single base pair substitution with MAF ≥ 0.8, [2] there were at least 15 viable matched background samples, and [3] there was at least one CpG site within 10 kb of the mutated base. Filtering against these criteria left 2,600 mutation events. Random control events were chosen to create a background distribution of ΔMF values at genomic loci lacking somatic mutations in any sample. For each true mutation event, we randomly chose 100 non-mutated nucleotides from the corresponding mutated sample and calculated ΔMF at these loci. To perform this calculation, we treated the randomly chosen nucleotide as if it were a mutation and calculated the ΔMF of CpG sites within ±10 kb.

### Extent of mutation-associated methylation remodeling

To investigate the extent of methylation remodeling associated with somatic mutation, we focused on the 25% of mutation events with the most positive or negative ΔMF values (**Fig. 4a**). We extended the range of ΔMF calculation to all measured CpG sites within ±100 kb from each site of mutation, computing ΔMF at overlapping 2 kb windows across this range. Then we aligned these these ΔMF values by their linear genomic distance from their respective mutated site (**Fig. 4c**).

### CpG and CGI enrichment

The genomic background rate of CpG islands (CGIs) and CpGs was calculated based on the hg19 annotation^95,96^ (https://hgdownload.soe.ucsc.edu/goldenPath/hg19/database/cpgIslandExt.txt.gz). The frequency of CpGs in CGIs was based on previously published statistics^50^. To understand wether mutation type (CpG or non-CpG) and location (CGI or non-CGI) were related to the degree of mutation-associated methylation change, we divided the frequency of each mutation type by the expected genome-wide rate (‘fold enrichment’, focusing on the 25% of mutation events with the most positive or negative ΔMFs). A one-sided binomial statistic was used to test for an increase in mutation frequency above the genomic background rate of each mutation type (**Fig. 4d**).

### Clock datasets and features

Tissue samples used for all clock-related analyses were from the LGG (Brain), GBM (Brain-2), SARC (Bone), KIRP (Kidney), or THYM (Thymus) cancer types in TCGA (n = 1,250 individuals). Somatic mutations within the 25 genes having the highest mutation frequency for that tissue type were removed to mitigate the influence of driving cancer genes on the analysis. We created a shared feature set for training all clock models, selecting the 100,000 CpG sites with the greatest average somatic mutation burden across samples within ±10 kb of the CpG site.

### Mutation clock

The mutation clock was based on a gradient boosted tree model, an XGBoost Regressor^97^ with default parameters, which we trained to predict chronological age using features derived from somatic mutations at the 100,000 CpG sites described above. In particular, the features used to describe an individual sample were (1) the counts of all somatic mutations within ± 10 kb of each CpG site (the “mutation burden”, **Fig. 5a**); (2) the one-hot encoded tissue type; and (3) the genome-wide mutation burden, summing the mutated allele frequencies (MAFs) across all 100,000 CpG sites. The accuracy of age prediction was assessed using nested cross validation, where 64% of samples were used for model training, 16% for hyperparameter tuning, and 20% for testing, with this entire procedure repeated over five folds (**Fig. 5b, Supplementary Fig. 2**). Following hyperparameter tuning, the number of features selected for use in the trained mutation clock model ranged from 1,711 to 1,781 across folds, with a mean of 1,759.

### Methylation clock

The methylation clock was also based on an XGBoost Regressor model, with identical default parameters^97^ as per the mutation clock described above. The definition of features was also closely matched, focusing on the same 100,000 CpG sites but using methylation rather than mutation data. In particular, features used to describe an individual sample were (1) the methylation fraction of each CpG; (2) the one-hot encoded tissue type; and (3) the overall degree of genome-wide methylation, computed as the sum of methylation fraction values across all 100,000 CpG sites (**Fig. 5a,c**, **Supplementary Fig. 2**). Nested cross validation was performed as for the mutation clock. Following hyperparameter tuning within each fold of cross validation, the number of features selected for use in the trained model ranged from 4,741 to 4,824 across cross-validation folds, with a mean of 4,787.

### Application of existing clocks

The Hannum, Horvath, and PhenoAge clocks were refit to the TCGA pan-cancer dataset, following a previously described procedure^33^. Briefly, the CpGs used in these clocks were obtained from the respective publications^33,34,39^ and a linear regression model^98^ with default parameters was trained on these features to predict the chronological age of TCGA samples. Some CpG sites included in these clocks did not have methylation values passing quality controls (see above Methods section “Data access and preprocessing”), so only the remaining 72%, 69%, and 76% of features were used in the re-fit Hannum, Horvath, and PhenoAge clocks, respectively. Nested five-fold cross validation was used to assess the performance of each of these re-fit clocks (**Supplementary Fig. 3a-b**). For each sample, the residual of each methylation clock (predicted age – chronological age) was compared with the residual of the mutation clock (**Fig. 5e**).

### Local association of methylation, mutation burden, and age

Across the 1,250 individuals and 100,000 CpG loci used in the mutation and methylation clocks, we compared the association between the methylation fraction of each CpG site, its mutation burden in the surrounding 20 kb, and the chronological age of the samples (**Fig. 5f**). First, we calculated the mutual information (MI)^99^ between the methylation fraction and mutation burden (±10 kb) at each CpG locus. Second, we selected the 1%, 5%, and 10% of CpG sites with the largest methylation-age MI and mutation burden-age MI and counted the number of CpGs shared between these groups. Third, we compared this overlap to the expected rate of overlap assuming random selection from the 100,000 original CpGs. A two-sided binomial test was applied to assess statistical significance.

### Software

All analyses were performed in the Python 3.10 and R 3.6.1 environments. Data analysis was conducted using Pandas 1.5.3, SciPy 1.10.0, and Statsmodels 0.13.5. Data were visualized with Seaborn 0.12.1 and Matplotlib 3.7.1.

## Acknowledgements

This study was funded by the National Institutes of Health under awards U54 CA274502 and P41 GM103504.

## Author Contributions

ZK designed the study, carried out the primary data analyses, and wrote the manuscript. AL and DE assisted with data analysis and study design considerations. TI and SC designed the study and wrote the manuscript.

## Declaration of Interests

T.I. is a cofounder of Serinus and Data4Cure, is on their Scientific Advisory Boards, and has equity interest in both companies. T.I. is on the Scientific Advisory Board of Ideaya BioSciences and has an equity interest. The terms of these arrangements have been reviewed and approved by the University of California San Diego, in accordance with its conflict of interest policies.

## Supplementary Figures

**Supplementary Figure 1:**
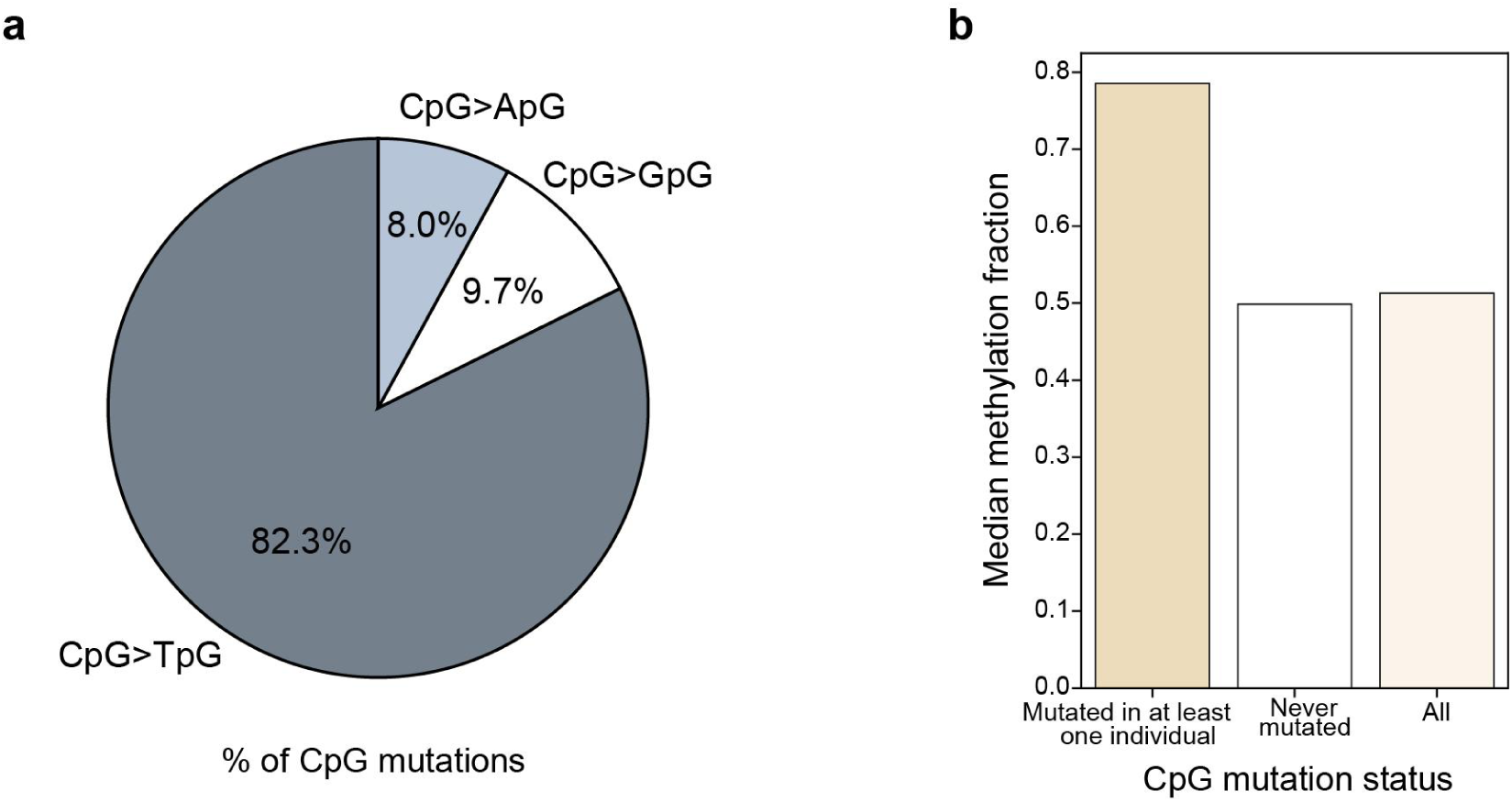
Supplemental characterization of CpG mutations. **a)** Pie chart showing the proportion of CpG mutations (n = 467,079 mutations) that result in specific mutated nucleotides. Note that 5’-CpG-3’ sites are palindromic, corresponding to a 3’-GpC-5’ sequence on the opposite strand; thus, mutation of the C residue is equivalent to mutation of the complementary G residue. For simplicity, we refer to all CpG mutations by the status of the C residue. **b)** Barchart showing the median methylation fraction across all PCAWG samples, considering CpG sites where a mutation has occurred in at least one sample (left, n = 1,137 CpG sites), CpG sites where no mutation has occurred in any sample (middle, n = 325,614 CpG sites), and all measured CpG sites (right, n = 326,751).

**Supplementary Figure 2:**
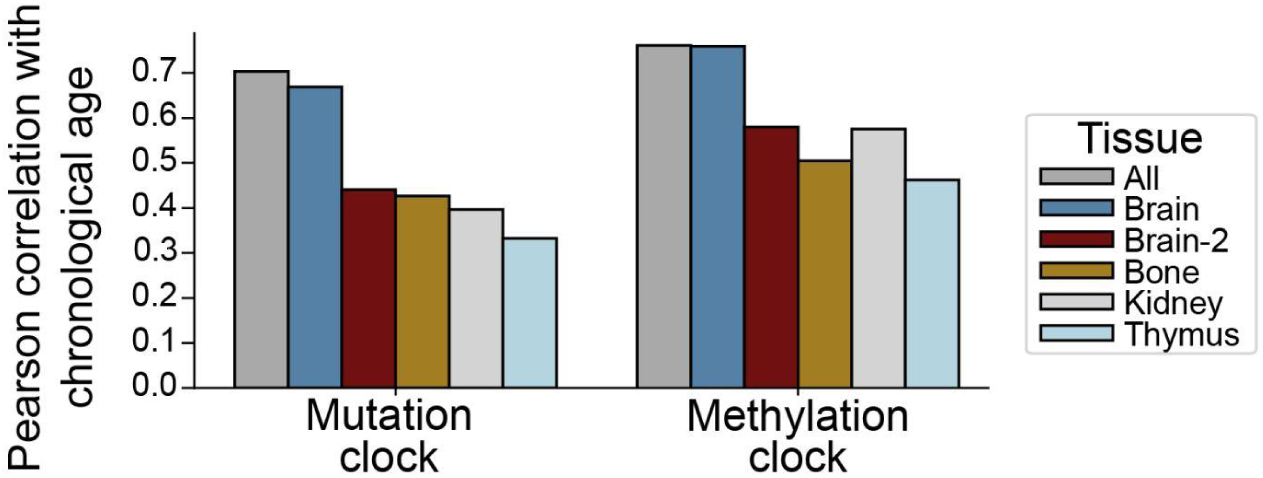
Supplemental age prediction accuracy. Bar plot indicating the correlation of chronological age with the age predictions of mutation versus methylation clocks across individuals (n = 1,250). Correlations are shown across all tissues and in each of five TCGA tissues individually: LGG (Brain), GBM (Brain-2), SARC (Bone), KIRP (Kidney), and THCA (Thymus).

**Supplementary Figure 3:**
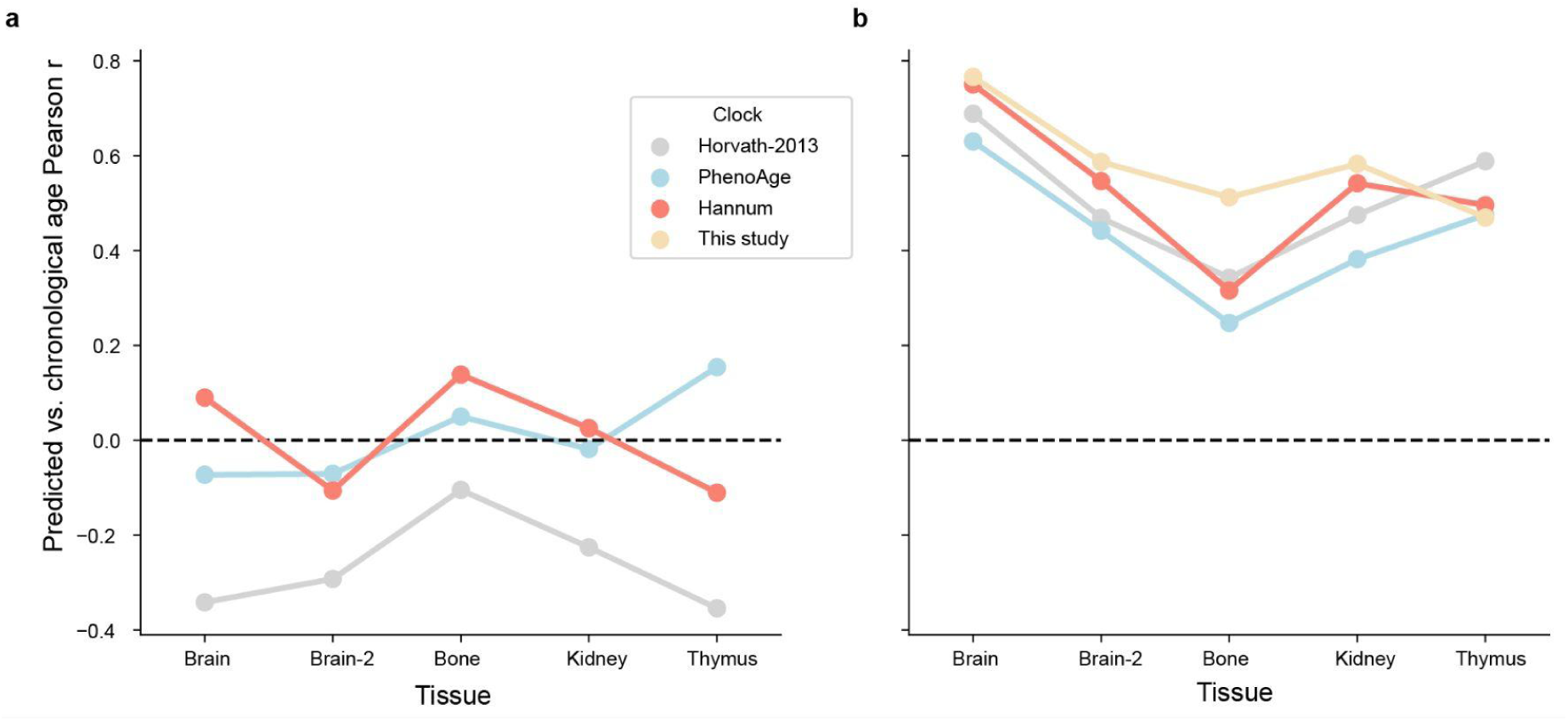
Performance comparison to previous epigenetic clocks. **a)** Pearson r between predicted and chronological age for Hannum, Horvath, and PhenoAge clocks across the same samples as Fig. 5b (n = 1,250). Nested five-fold cross validation. The performance of the methylation clock trained in this study (“This study”) is shown for reference. **b)** Pearson r between predicted and chronological age for Hannum, Horvath, and PhenoAge clocks after re-fitting (**Methods**). Same samples and validation procedure as (a).

